# Centromeric repeats of the Western European house mouse I: high sequence diversity among monomers at local and global spatial scales

**DOI:** 10.1101/2020.08.28.272245

**Authors:** William R. Rice

## Abstract

Previous work found that the centromeric repeats of the Western European house mouse (*Mus musculus domesticus*) are composed predominantly of a 120 bp monomer that is shared by the X and autosomes. Polymorphism in length and sequence was also reported. Here I quantified the length and sequence polymorphism of the centromeric repeats found on the X and autosomes. The levels of local and global sequence variation were also compared. I found three length variants: a 64mer, 112mer and 120mer with relative frequencies of 2.4%, 8.6%, and 89%, respectively. There was substantial sequence variation within all three length variants with a rank-order of: 64mer < 120mer < 112mer. The 64mer was never found alone on long Sanger traces, and was arranged predominantly as a 176 bp higher-order repeat composed of a 64/112mer dimer. Reanalysis of archived ChIP-seq reads found that all three length variants were enriched with the foundational centromere protein CENP-A, but the enrichment was far higher for the 120mer. This pattern indicates that only the 120mer contributes substantially to the functional centromeres, i.e., to the kinetochore-binding, centric cores of the centromeric repeat arrays. Despite only moderate sequence divergence among random pairs of 120mers (averaging 5.9%), other measures of sequence diversity were exceptionally high: i) variant richness (numerical diversity) –on average, one new sequence variant was observed every 4th additional monomer randomly sampled (in N = 7.2 × 10^3^ monomers), and ii) variant evenness –all of the nearly 2 × 10^3^ observed sequence variants were at low frequency, with the most common variant having a frequency of only 5.7%. I next used long Sanger trace data from the Mouse Genome Project to assess the pattern of monomer diversity among neighboring 120mers. Unexpectedly, side-by-side monomers were rarely identical in sequence, and sequence divergence between these neighbors was nearly as high as that between random pairs taken from the genome-wide pool of all 120mers. I also used long Sanger traces to determine sequence variation among neighborhoods of 5 contiguous 120 bp monomers. Sequence diversity within these small regions typically spanned most of the entire range of that found genome-wide. Despite high sequence variation within these neighborhoods, the density of monomers with functional binding motifs for CENP-B (i.e., b-boxes with sequence NTTCGNNNNANNCGGGN) was strongly conserved at about 50%. The overarching pattern of monomer structure at the centromeric repeats of this subspecies is: i) high homogeneity in the density CENP-B binding sites, and ii) high heterogeneity in monomer sequence at both local and global levels.

## Introduction

Centromeres are the regions of DNA that assemble a kinetochore and thereby attach a chromosome to the microtubules of the mitotic or meiotic spindle (Mitchell 1996; Westhorpe and Straight 2015). Across a broad range of species, centromeric DNA shows little sequence conservation, even among closely related species (Henikoff et al. 2001) –although centromeres typically dosh are the more general characteristics of being composed of long tandem repeat arrays with higher than average A/T content (Müller and Almouzni 2017). One exception to this lack of sequence conservation in some species is a 17 bp binding site for **CEN**tromere **P**rotein **B**(CENP-B; Masumoto et al. 1989; Fachinetti et al. 2015). I will refer to this binding site as a ‘b-box,’ with consensus sequence 5’-CTTCGTTGGAAACGGGA-3’ in humans (Masumoto et al. 1989). Available evidence indicates that CENP-B is present in most mammals (Yoda et al. 1992; Casola et al. 2008), yet in most of these species it does not bind at centromeres (Haaf et al. 1995; Alkan et al. 2011; Kugou et al. 2016) –presumably because a functional b-box binding sequence (NTTCGNNNNANNCGGGN; Masumoto et al. 1989; Kipling et al. 1995; Kipling and Warburton 1997) is absent.

Both the Western European house mouse (*Mus musculus domesticus,* and many other species within the genus *Mus)* and humans (and all other great apes) are unusual among mammals because they bind CENP-B at the centromeres of nearly all of their chromosomes –only the Y chromosome lacks this binding. Despite sharing this unusual trait, the organization of the centromeric repeats of these species are highly dissimilar.

Human chromosomes are primarily metacentric with a minority of acrocentrics and no telocentrics. Their centromeric repeats on the X and autosomes are organized into **H**igher **O**rder **R**epeats (HORs – repeats of sets of 2 or more monomer types). These tandemly repeated units contain 2 to 19 different monomers (each ~170 bp; see Rice 2019A for a complete listing of the consensus sequences of all 24 human centromeres from a singles genome). Monomers can differ in sequence within an HOR by as much as 20%, and between HORs by as much as 35% (Willard 1991; Rice 2019A). Monomers within human HORs are predominantly organized into dimers (Masumoto et al. 1989; Thomas et al. 1989), with one monomer containing a 17 bp b-box that binds CENP-B, and the other containing a 19 bp ‘n-box’ (consensus = T[G/A][G/ A]AAAAGGAAATATCTTC; Rice 2019A).

In sharp contrast, the X and autosomes of the Western European house mouse are all telocentric (in most laboratory strains and most wild populations – and in contrast to the acrocentric Y chromosome). Their centromeric repeat arrays (minor satellite) are reported to lack HORs (Broccoli et al. 1990; 1991; Kipling et al. 1994; Pertile et al. 2009; Komissarov et al. 2011) and contain a 120 bp monomer (Wong and Rattner 1988) that has much less sequence variation than human centromeric monomers (Kipling et al. 1994). Despite the lack of HOR structure, evidence for substantial monomer variation in both length and sequence was reported soon after mouse centromeric monomers were first discovered (e.g., Broccoli et al. 1991; Kipling et al. 1994). A recent study by Iwata-Otsubo et al. (2017) used whole genome shotgun (WGS) sequencing to quantify sequence variation among the 120 bp centromeric monomers. They found six polymorphic SNPs with exceptionally high variation, eleven additional SNPs with high polymorphism, and that less than half the 120 positions had little or no variation. Note that I use the term ‘polymorphism’ here to denote base pair variation among aligned centromeric monomers irrespective of their genomic location.

Centromeric repeats differ substantially in sequence among species within the genus *Mus* (Wong et al. 1990). These repeats also vary substantially in array size (as measured by RFLP polymorphism) between different strains of mice–including a pair of congenic strains that were separated by only 20 years (Aker and Huang 1996). These observations suggest that mouse centromeric repeats evolve rapidly within and between species, as has been reported in many other diverse taxa (Henikoff et al. 2001).

Here I characterize and quantify the variation of centromeric monomers within the Western European house mouse. I do this by analyzing archived long WGS Illumina reads (150 bp) from strain C57BL/6J (Wang et al. 2018) and longer Sanger traces (up to ~1 kb) generated during the Mouse Genome Project on this same mouse strain (Waterson et al. 2002). I found exceptionally high levels of numerical diversity (high richness or number of sequence variants) and that no single sequence variant was common (high evenness –the relative frequency of the most common variant was < 6%) despite only moderate sequence divergence among monomers (averaging 5.9%). I also found that matching sequences were rare among neighboring monomers and that local monomer sequence diversity was nearly as high as global diversity. I hypothesize that this unusual repeat structure has evolved in response to selection for resistance to deletion after repair of double strand breaks (DSBs) via the **S**ingle **S**trand **A**nnealing (SSA) pathway. In the companion paper I expand on this hypothesis and explore the possibility that centromere position contributes to coevolution between centromeres size (bp) and karyotype number (2N).

## Methods

Archived data bases of WGS sequences were searched for reads containing centromeric repeats using either BLASTN (**B**asic **L**ocal **A**lignment **S**earch **T**ool-**N**ucleotide) or a sliding window to search for sequences closely matching the consensus 17 bp CENP-B-binding sequence (which in *M . m . domesticus* is ATTCGTTGGAAACGGGA = consensus b-box; Iwata-Otsubo et al. 2017). Most analyses were carried out using large samples from two archived data sets: i) paired-end illumina reads (150 bp) from WGS sequencing of the genome of the Western European house mouse strain C57BL/6J (SRA file SRR6339177; available at url https://trace.ncbi.nlm.nih.gov/Traces/sra/?run=SRR6339177) described in Wang et al. (2018), and ii) Sanger traces from the Mouse Genome Project (strain C57BL/6J; available at url ftp://ftp.ncbi.nih.gov/pub/TraceDB/mus_musculus/) described in Waterson et al. (2002). Another archived data set was used to a more limited degree to evaluate centromeric activity of different monomer length variants. In this case, I used large samples from archived ChIP/Input WGS Illumina reads (100 bp) described in Iwata-Otsubo et al. (2017). The Input data file was SRR 5723792 (https://trace.ncbi.nlm.nih.gov/Traces/sra/?run=SRR5723792) and the ChIP file (using antibodies against the centromeric protein CENP-A) was SRR 5723794 (https://trace.ncbi.nlm.nih.gov/Traces/sra/?run=SRR5723794).

## Results

### Monomer length variants

Because published FISH studies found CENP-B to be highly concentrated at all of the centromeres of the X and autosomes within the genome of *M. m. domesticus* (and nowhere else, e.g., Kipling et al. 1995; Fachinetti et al. 2015), I first screened a large sample (N = 19,977,056) of Sanger traces from the Mouse Genome Project (trace files 1-49) for b-boxes (CENP-B binding sites), and when found, I extracted their sequences and the intervening sequence between them. The search was performed by moving a 17 bp sliding window along a trace and recording the position of each window that sufficiently matched the b-box sequence (or the complementary sequence on the opposite strand of DNA). The sufficiency of a match to the b-box sequence was determined by previously published polymorphism data for this sequence in strain C57BL/6N.

Polymorphism data from a recent study (Iwata-Otsubo et al. 2017) indicated the the b-box sequence of Western European house mouse 120 bp monomers is highly polymorphic (by which I mean sequence variation between b-boxes within monomers from all genomic locations – within and between chromosomes) with 3 positions containing very high levels of polymorphism and 4 additional positions with lower but substantial levels of polymorphism (see Supplemental Figures S1 and S2 of Iwata-Otsubo et al. 2017). For this reason, I scanned (with a sliding window) the Sanger traces for any sequences that differed by ≤ 4 bp from the 17 bp consensus b-box sequence. I also included 7 b-box sequences that deviated by 5 bp that, by trial-and-error (i.e., including sequences in the sliding window analysis that differed by > 4 bp), I found to be present at non-trivial frequency in mouse centromeric satellites. The 50 most common b-box sequences (and their frequency of occurrence) are shown in Supplemental Figure S1. Every time two successive matches were found, I recorded the starting and ending presumptive b-box sequence and also the intervening sequence. I defined a centromeric unit to be the sequence of the first b-box and the following intervening sequence before the next b-box.

A histogram of the length of centromeric units revealed three well-defined peaks at 64, 112, and 120 bp (Supplemental Figure S2). I next excluded any units that did not start (in either sequencing direction) before the final 137 bp of each trace so as to not bias the search toward the shorter 64 and 112 bp centromeric units, i.e., I ensured that there was sufficient space to find all three monomer length variants. As a check on these results, and because all three centromeric units were ≤ 120 bp, I also ran a sliding window analysis on my sample of 150 bp Illumina reads from strain C57BL/6J (SRR6339177, Wang et al. 2018). The proportion of centromeric units of sizes 64, 112, and 120 bp (each ± 2 bp) are shown in Figure 1.

**Figure 1.**
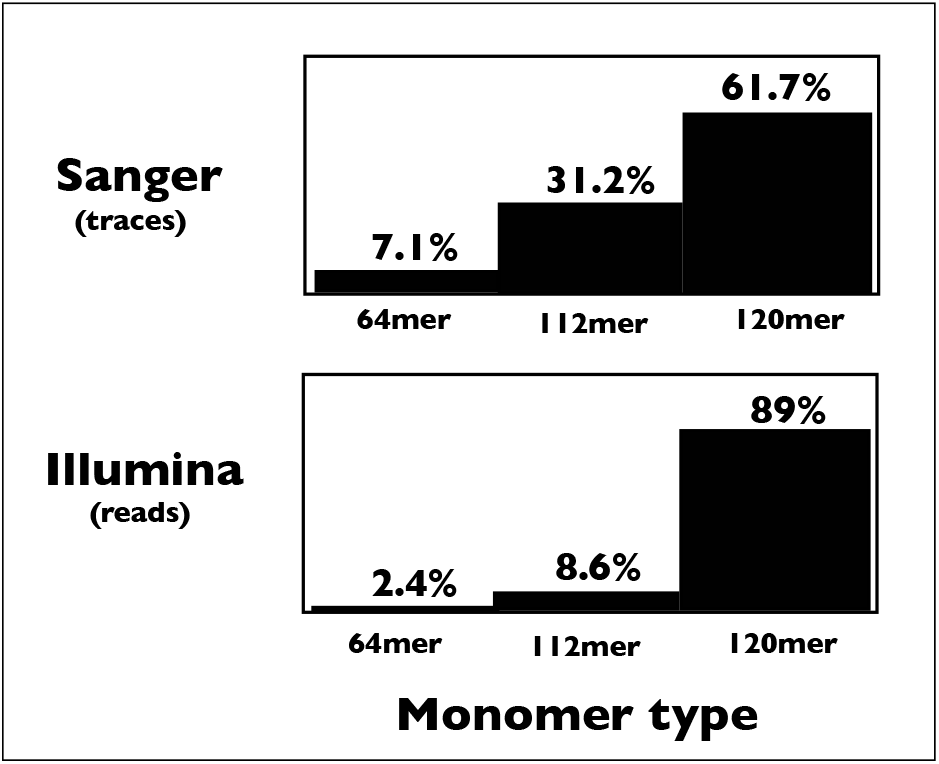
Histograms of the proportion of the three monomer types (size classes) in the sample of Sanger traces (variable length) and Illumina 150 bp reads.

Unexpectedly, there was strong disagreement in the frequencies of the three size units between the Sanger and Illumina data sets –with the 120mer being far less common in the Sanger traces. This discrepancy could feasibly occur if the cloning step used in Sanger sequencing selected against the 120 bp units. To test for this possibility, I used logistic regression to examine size category frequencies vs. trace length. I found a very strong reduction in the frequency of the 120 bp centromeric unit as trace length increased (Supplemental Figure S3) which could feasibly account for the much lower frequency of the 120 bp units in the Sanger compared to the Illumina data sets. For this reason I concluded that the estimates of the frequencies of the three size categories of centromeric monomers from Sanger tracers were unreliable, and used the Illumina data to estimate these frequencies in the Western European house mouse genome (strain C57BL/ 6J).

Although the longer Sanger trace data set from the Mouse Genome Project was biased against long traces containing 120 mers, I was nonetheless able to use this data set to examine how the three units are packaged in centromeric repeat arrays. To visually examine the spatial arrangement of neighboring monomers, I examined a sample of 608 long Sanger traces that contained at least 5 monomers each. Despite the 64mer unit being more prevalent in longer traces, the longest stretch of tandem 64mers that I observed was only three in a row and this run-length was extremely rare (only one case observed). Instead, 64mers were arranged predominantly as a 176 bp dimer containing one 64mer and one 112mer (Figure 2, Top). Most 112mers and 120mers were arranged as tandem repeats of the same length monomers (Figure 2, middle). In the rarer cases where both 112 and 120 monomers were present within a trace, one or the other predominated (Figure 2, bottom). Because the 64, 112 and 120 bp centromeric units were arranged as tandem repeats (as a dimer in the case of the 64 bp unit), hereafter I will refer to them as centromeric monomers.

**Figure 2.**
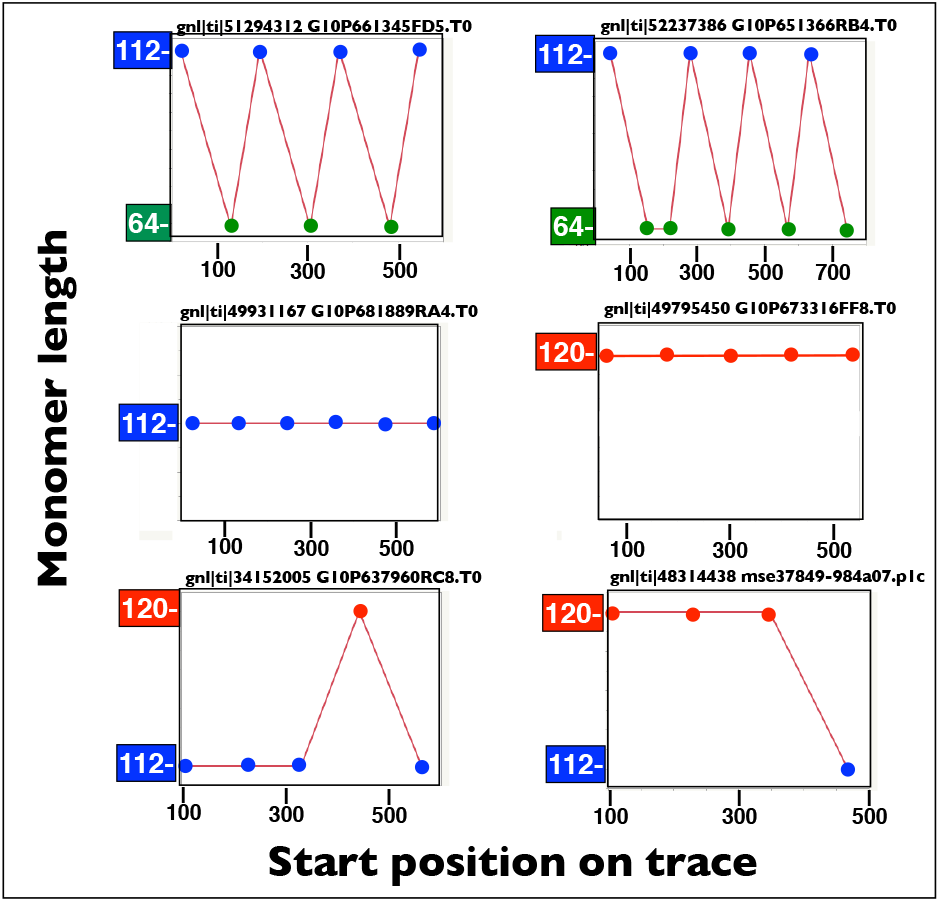
Representative examples of the organization of contiguous monomers in long Sanger traces. **Top row.** The 64mers were most commonly observed as runs of 64mer/112mer dimers (top left) with occasional tandem repeats of one monomer type (top right). **Middle and Bottom rows.** The 112mer and 120mers monomers were most commonly observed as monotypic runs of one or the other monomer type (Middle) with infrequent cases in which one monomer did not match the other monomers (Bottom).

Centromeric monomers are expected to diverge in sequence via mutation when they are isolated by distance within a single centromeric array and/ or when they are located on different chromosomes (Smith 1976; Roizès 2006). Unequal crossing over (Smith 1976) and also out-of-register **B**reak-**I**nduced **Repair** (BIR; Rice 2019B) within a repeat array will act to homogenize nearby sequences. Ectopic exchange –as might occur, for example, via out-of-register gene conversion within or between repeat arrays– will act to homogenize more distant monomer sequences. All else being equal, monomers that originated more recently are expected to have less standing genetic variation because there has been less time for mutations to accumulate among copies. I compared the variation in sequence among the monomer size types. Because the 64mer is much shorter, it is expected to be less variable on a unit-wide basis.

The consensus sequence of the three monomer size classes are shown in Figure 3. To compare levels of sequence variation among the three monomer types, I first compared the level of sequence variation among the 1^st^ 64 bp of each type –which are predominantly homologous (Figure 1: note this region [left half in Figure 1] contains the b-box region that is a hot-spot for polymorphism in the 120mer; see Supplemental Figures S1 and S2 in Iwata-Otsubo et al. 2017). To minimize artifacts from sequencing error, here and in all remaining analyses of sequence divergence (unless otherwise indicated), I only used reads in which all base calls had an accuracy of ≥ 99.9%. The 64mer was estimated to have 28.8% less sequence variation compared to the 112mer and 11.8% less variation compared to the 120 mer (Figure 4, red bars, P < 0.001 for both comparisons with nonparametric median test). To compare the 120mer and 112mer, I used their entire sequence. Although slightly longer, the 120mer was estimated to have 25.3% less sequence variation compared to the 112mer (Figure 4, blue bars, P < 0.0001 with a nonparametric median test). So in summary, the ordering of genetic variation is 112mer > 120mer > 64mer.

**Figure 3.**
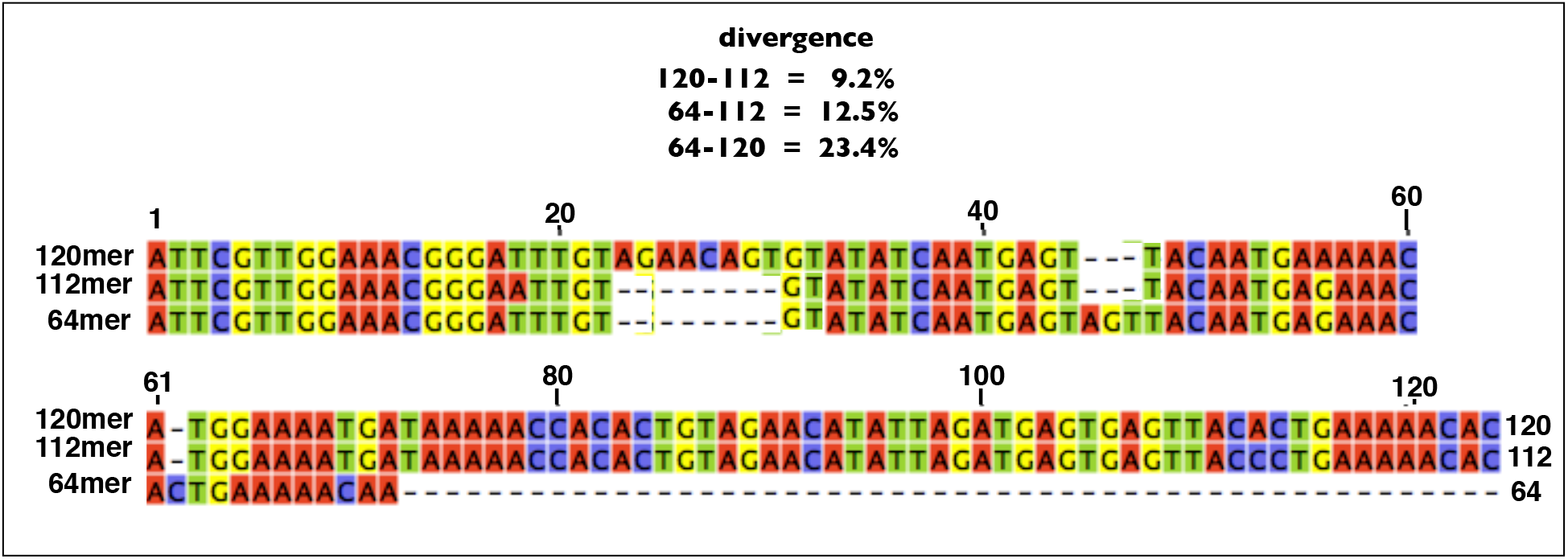
Alignment of the consensus sequences for the 64mer, 112mer and 120mer monomers. Divergence from the 64mer excludes its terminal 48 bp deletion.

**Figure 4.**
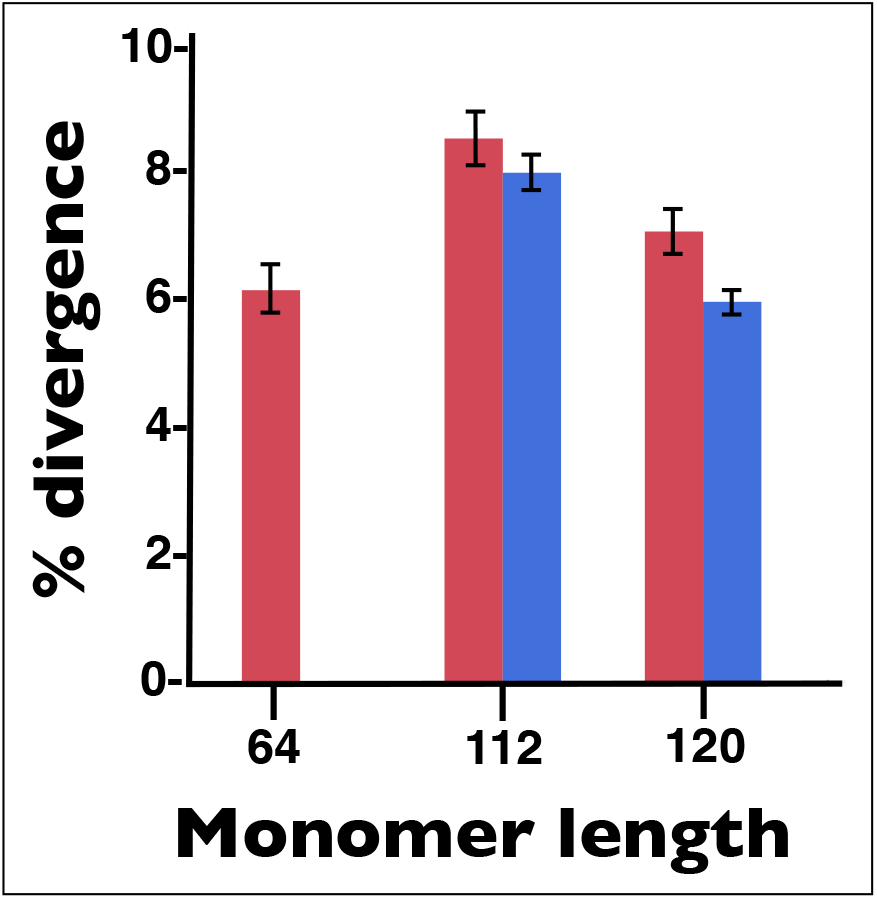
Histograms of the average percent divergence between randomly sampled monomers of the same length category. The red bars represent divergence among the 1st 64 bp of each size category and the blue bars include all positions in the 112mer and 120mer. Error bars represent 95% confidence intervals. Sample sizes per histogram bar were N = 500 aligned pairs of sequences.

The three monomer length variants differed substantially in the frequency of mutated b-boxes that are predicted to not bind CENP-B, i.e. those that deviate from the minimal sequence required to bind CENP-B: NTTCGNNNNANNCGGGN) (Masumoto et al. 1989; Kipling et al. 1995; Kipling and Warburton 1997). I will refer to b-boxes that are predicted to bind CENP-B as “functional” and those that do not as “non-functional.” The percentages of functional b-boxes were 72, 45, and 41 for the 64mer, 112mer and 120mer respectively (Supplemental Figure S4).

I next determined the contribution of the three size classes of centromeric monomers to centromere functioning. Kinetochores are assembled over a subset of a centromeric repeat array (the centric core) that is strongly enriched with the histone variant **CEN**tromere **P**rotein-**A**(CENP-A; Palmer et al. 1987, 1991). For this reason, ChIP-seq data using antibodies against CENP-A can be used to determine the monomeric types that are currently being used as the functional centromere (e.g., see Bodor et al. 2014). This type of data was generated by Iwata-Otsubo et al. (2017) when analyzing the 120mer monomer, but their data base can be reanalyzed to look at all three of the predominant monomer size classes described here. This data set indicates that all three monomer size classes have elevated ChIP/Input ratios compared to a major satellite control (Figure 5), which was selected because it has similar GC content, similar genomic location, and shares long stretches of sequences with as high as 83% sequence identity (Wong and Rattner 1988). Although the ChIP/Input ratio was elevated relative to control for all three monomer size classes, it was far higher for the 120mer monomers (Figure 5). This pattern indicates that the 64mer and 112mer size classes most plausibly reside near –but outside– the centric core region of the repeat arrays that recruit the kinetochore on the X and autosomes. For this reason, I next focus exclusively on the 120mer.

**Figure 5.**
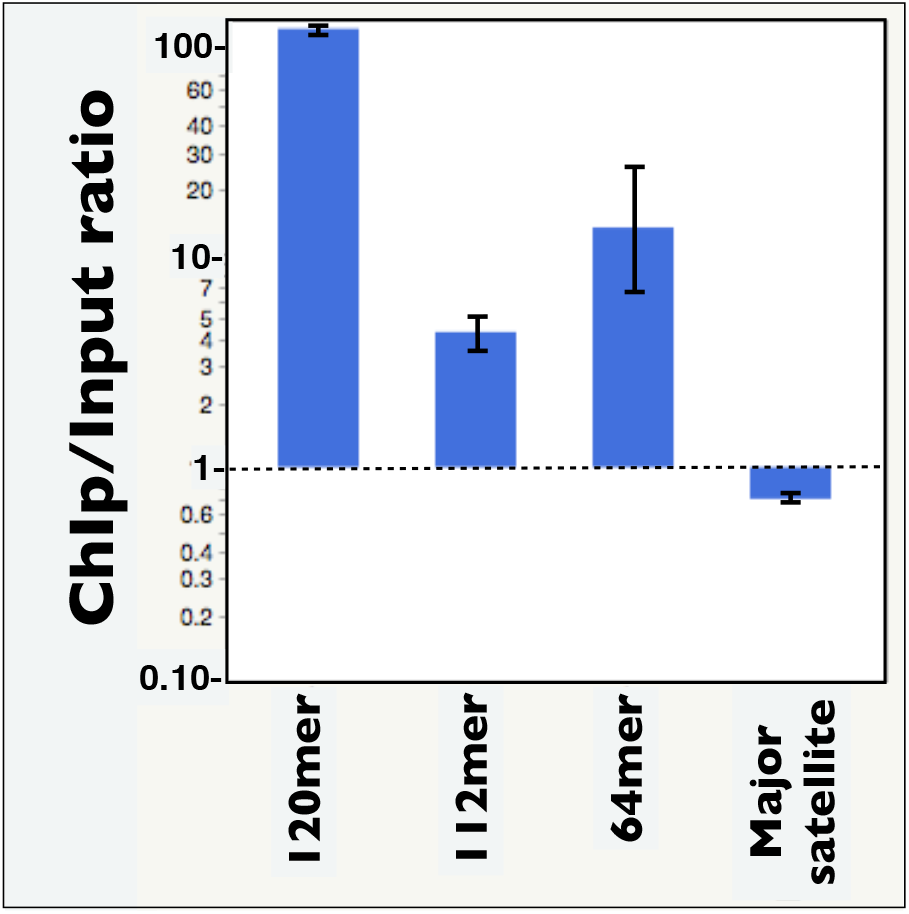
Ratios of the read depth of monomer sequences (from BLASTN) from equal sized Input and ChIP samples (each with N = 1.5 × 10^6^ reads). Error bars represent 95 bootstrap confidence intervals.

### Monomer diversity of the 120mer

To characterize the sequence variation of the 120mers, I began with a sample of 7,218 120mers (obtained using a sliding window screen of 150 bp Illumina reads from C57BL/6J; from SRA file SRR6339177; Wang et al. 2018) that had all base calls with an accuracy of ≥ 99.9%. I grouped this random sample of monomer sequences with low sequencing error into 1,923 bins of identical sequence and then rank-ordered the sequence bins from most to least common. I then made a neighbor-joining cluster diagram of the 100 most common sequence bins (Figure 6). The 120mers have two major sequence clusters: one with all members with nonfunctional b-boxes (blue) and the other with predominantly functional b-boxes (red). The cluster with non-functional b-boxes has a greater degree of sequence variation, i.e., it has substantially wider spread in Figure 6.

**Figure 6.**
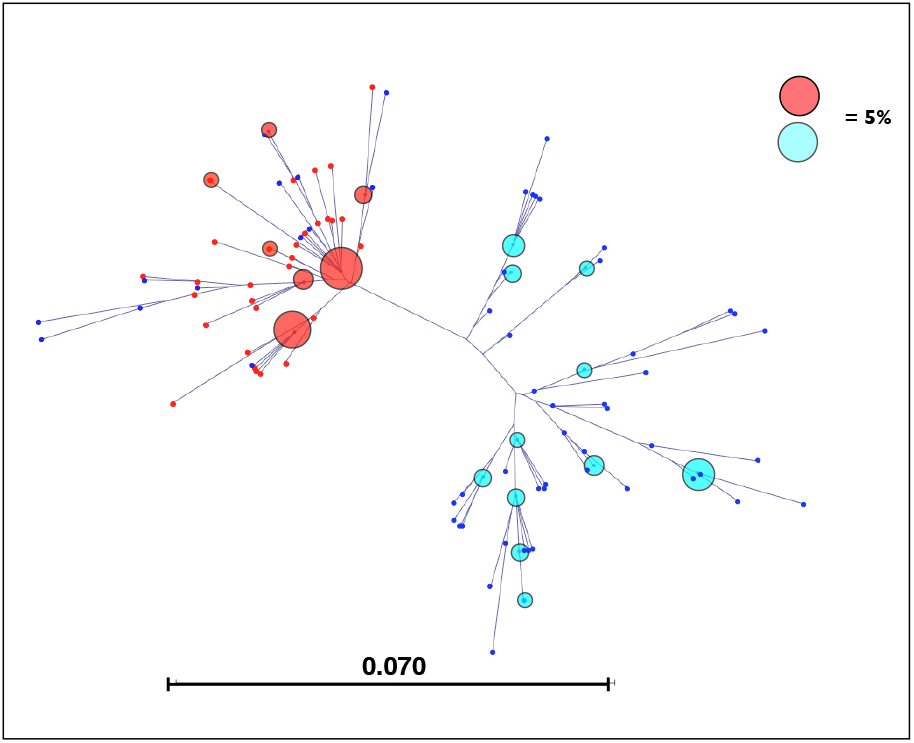
A cluster diagram (neighbor-joining tree) of the 100 most common 120mer sequences. Red nodes contain functional b-boxes (predicted to bind CENP-B) and blue nodes contain non-functional b-boxes (predicted to not bind CENP-B). All nodes with a frequency > 0.6% are shown with an enlarged, semi-transparent blue or red circles, the size of which is scaled relative to a 5% benchmark.

Average sequence divergence f rom the consensus 120mer was estimated from a sample of low-error (all base calls had accuracy ≥ 99.9%) Illumina sequences. Mean divergence was 3.73% (SD = 2.12%; N = 1000 aligned sequences) with 95% CI of (3.60, 3.86) and a maximum divergence of 10.2% (Supplemental Figure S5). The frequency spectra of monomer classes (i.e., a plot of sequence abundance vs. the rank of sequence abundance) had a total of 1,923 different sequences (from a sample of 7,218 low-error monomers [all base calls had accuracy ≥ 99.9%]), and was dominated by rare sequences (Figure 7). The most common sequence (the consensus sequence) had a frequency of only 5.7% and 80% of the sequences had a frequency of less than 1%. Lastly (and as described earlier in Figure 4), I measured sequence divergence among random pairs of monomers. They differed in sequence by an average of 5.9% (SD = 2.49) with a 95% CI of (5.70, 6.13) (see Figure 4).

**Figure 7.**
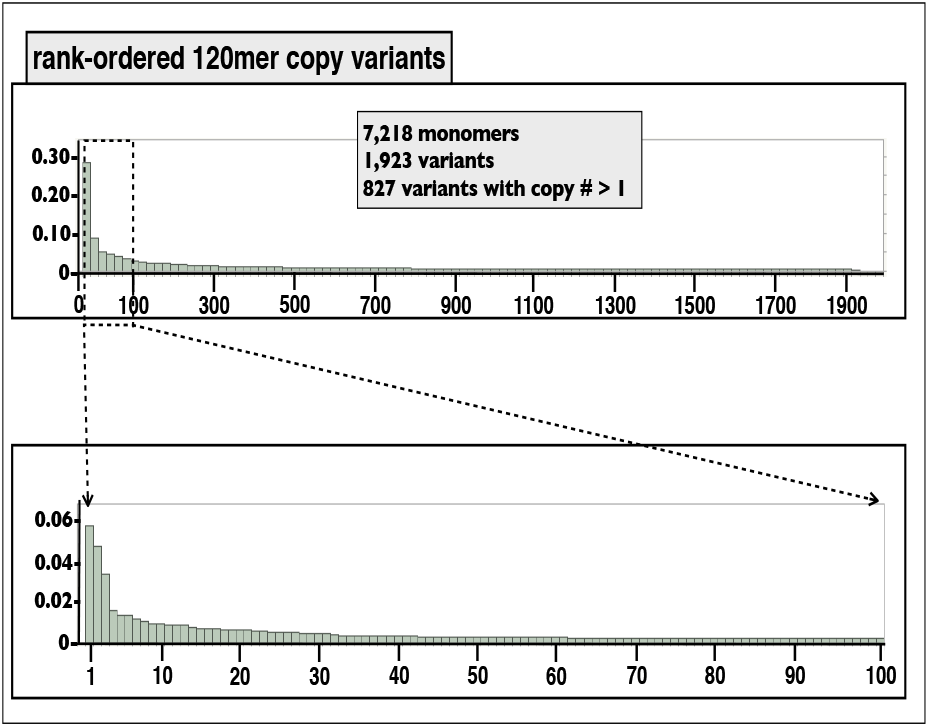
The frequency spectrum of 120mer sequence variants. The sample size was N = 7,218 random monomers, and the sequencing error rate was low (base call error rate of ≤ 0.001 at each of the 120 base pair positions). A total of 1,923 different sequences were observed and 827 of these had a frequency of at least two copies. No sequence was common (highest proportion was only 5.7%) and there was a long tail of many hundreds of sequences with similar frequency.

I next examined local sequence variation by analyzing patterns in Sanger traces that contain contiguous 120mers. I first examined a sample of N = 10,813 side-by-side 120mers that were sequenced with low error (*average* accuracy per base call ≥ 99.9%). Only 0.9% of the time were the neighboring monomers identical in sequence (Supplemental Figure S6), indicating that tandem copies of the same monomeric sequence are rare (95% CI [Wilson Score method] for occurrence rate = 0.48%, 1.74%). This value was similar to the proportion of time that randomly selected monomers from different Sanger traces were identical in sequence (0.8%, N = 2,082; Supplemental Figure S6).

Using the same low-sequencing-error sample of aligned monomers from the same traces, I next compared the level of sequence divergence between side-by-side monomers (N = 978) to those on different different traces (N = 1,481) and found the level of sequence divergence to be 93% as large between neighboring monomers (5.3%, 95% CI = 5.16%, 5.49%) as it was between monomers on different (random) traces (5.75%, 95% CI = 5.61%, 5.88%) (Supplemental Figure S7). A 95% conservative upper bound on this proportion (lower 95% bound for neighbors / upper 95% bound for non-neighbors) is 87.7%. So most of total sequence variation between random monomers is found between neighboring monomers –a surprisingly high level of diversity at the lowest possible level of spatial separation.

To expand my analysis of local sequence diversity beyond side-by-side monomers, I graphically examined a sample of 25 long Sanger traces that contained 5 contiguous 120mers. These represent a sample of ‘neighborhoods’ of 5 monomers. I next clustered each group of 5 contiguous monomers with a group of 14 reference monomers (Supplemental Figure S8) that span all the major clusters in the cluster diagram of all genome-wide monomers (Figure 6). These reference monomers represent the 13 most common monomers, genome-wide, plus the 18th most common monomer sequence, which was included because this addition generated a reference set that covered all the main branches of clustering of all 120mers (Figure 6). In Figure 8, I show the cluster diagrams for the 12 neighborhoods of 5 monomers with the highest sequencing accuracy and in Supplemental Figure S9 I show the remaining 13 clusters with lower sequencing accuracy. The pattern is the same in both subsamples: neighborhoods of only 5 contiguous monomers typically span most of the total, genome-wide sequence variation.

**Figure 8.**
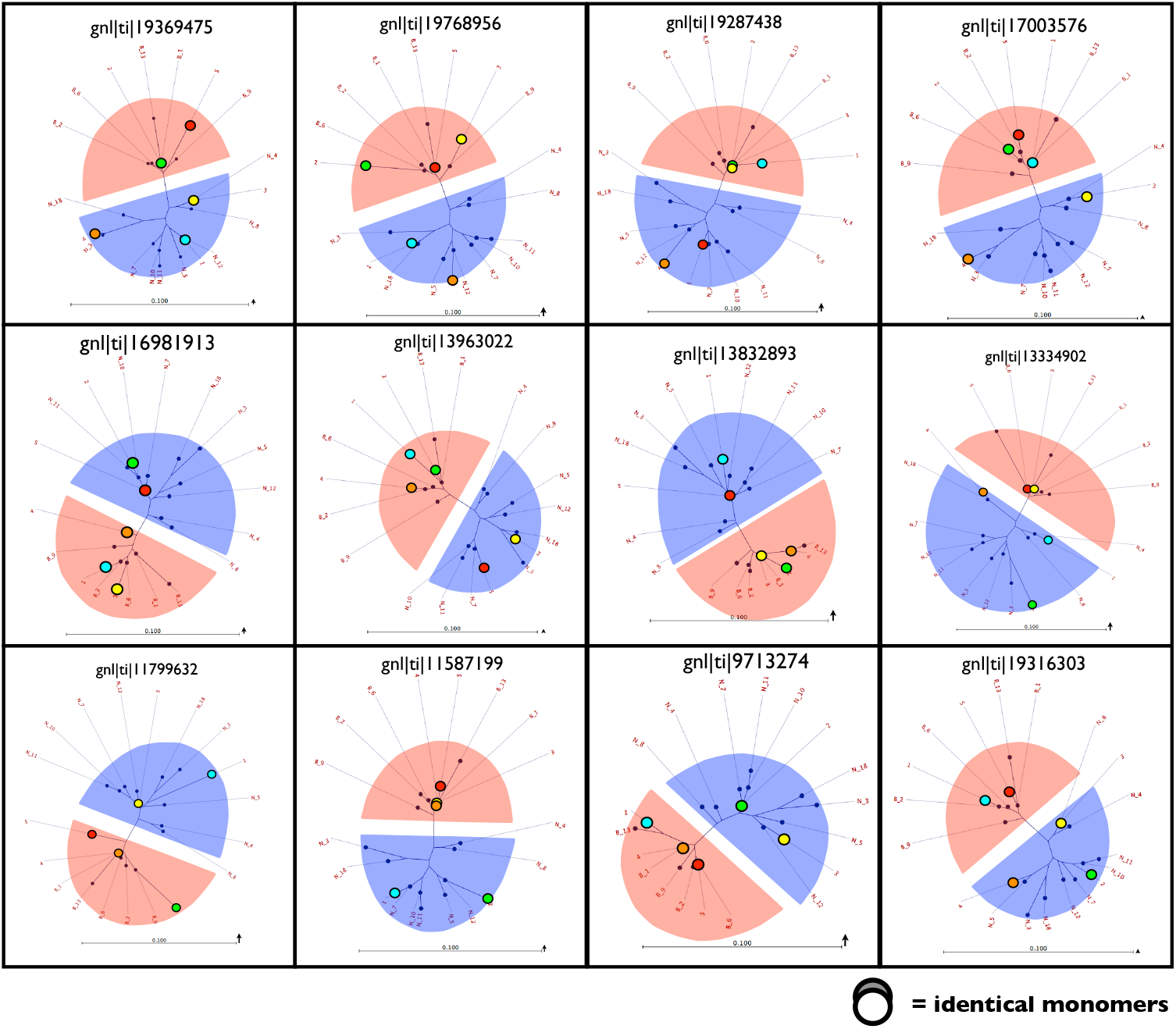
Cluster diagrams (neighbor-joining trees) of five contiguous 120mer monomers from long Sanger traces in combination with a reference group of monomers (14 total) containing the the most common 120mer sequences found genome-wide. The group of 14 represents the 13 most common sequences plus the 18th most common (added to have all the major branches of the cluster diagram shown in Figure 6 represented). The cluster diagram including only the reference groups is shown in Supplemental Figure S8). The orientation of each cluster diagram is determined by default parameters in the program CLC Sequence Viewer version 7.6.1, so the two major clusters (with and without functional 17 bp b-boxes that are predicted to bind CENP-B) are color coded: red with functional b-boxes and blue without functional b-boxes. Shown in the figure are the 12 Sanger traces with the lowest sequencing error rates. Average sequencing error across the five monomers is depicted by an arrow at the right edge of the scale bar at the bottom of each cluster diagram. The arrows depict the expected deviation (due to sequencing error) of each monomer sequence (contained in the 5 contiguous 120mers) from its position shown in the cluster diagram. The other 12 Sanger traces with higher error rates are shown in Supplemental Figure S9. Note that in both figures, a substantial amount of the sequence variation across the entire reference group is spanned by the 5 monomers from each Sanger trace. The order of the 5 neighboring monomers is color-coded: red (1^st^), orange, yellow, green, blue (last).

In humans, monomers (~170 bp) within HORs are configured predominantly as dimers: one monomer with and one without a functional b-box (see Figure 4 in Rice 2019A for the relative frequencies of these dimers across all human chromosomes). This configuration leads to a high degree of uniformity in the density of b-box- and no-b-box monomers. I noticed in my analysis of neighborhoods of 5 contiguous monomers in Sanger traces that most 5mers contained similar numbers of monomers with and without functional b-boxes (2-3 of each type). To test whether this pattern was likely by chance alone, I defined an ‘evenness’ statistic to be the abs([number of monomers with functional b-boxes] - [number of monomers with non-functional b-boxes]). This statistic is minimally one (high evenness) and maximally five (low evenness). Across the 25 5mers in Figure 8 and Supplemental Figure S9, the average evenness statistic was far smaller than predicted by chance (p = . 0003; Supplemental Figure S10). As a check on this pattern, I expanded the sample size to N = 57 5mers (all of those that I found in a sample of Sanger traces: recall that long Sanger tracers with 120mers were rare): the pattern persisted and the p-value declined to p = 0.00001 (Supplemental Figure S11). This pattern indicates that the density of b-boxes is far less variable than expected by chance with most 5mers having similar numbers of functional and non-functional b-box monomers.

## Discussion

Although the centromeric repeat sequence (monomer) of the Western European house mouse is commonly described as a 120 bp unit (Pietras et al. 1983; Wong and Rattner 1988), previous studies had also found two length polymorphisms: a 112 bp variant with an 8 bp deletion relative to the consensus 120mer, and a 64 bp variant that shares the 8 bp deletion in the 112mer but also had a large (48 bp) deletion that spans most of the 3‘ half of the consensus 120mer as shown in Figure 1 (see Kipling et al. 1994 for a compilation of many published minor satellite sequences). I found all three of these length length variants at non-trivial frequency (Figure 2) within Sanger traces from the Mouse Genome Project and also within long (150 bp) Illumina reads generated by Wang et al. (2018).

The 112mer has a frequency of about 9% (Figure 2B) and has the highest average pair-wise sequence variation among copies (8.47%; Figure 4). Based on my sample of Sanger traces, it is predominantly found as monotypic tandem repetitions (Figure 3). Its ChIP/Input ratio (~4 with antibody against CENP-A) is elevated compared to the major satellite control (Figure 5), but much lower than that of the more common 120mer (~114). This elevated-but-low ChIP/Input ratio indicates that the 112mer is probably in the region flanking the centric core of one or more of the the DNA repeat arrays that recruit the kinetochores of the X and autosomes (the centric core spans ~ 20% of the minor satellite repeat array on a typical telocentric chromosome; Iwata-Otsubo et al. 2017). The higher diversity of the 112mer, despite its smaller numbers and reduced length compared to the 120mer, is consistent with the hypothesis that this monomer is older and has been replaced by the less diverse 120mer as the predominant centromeric monomer.

The 64mer is the least common length variant (~2% of monomers) and it has the lowest average sequence divergence among monomer copies (Figure 4). No long repetitions of the 64mer were observed and a strong majority were arranged as 176 bp dimers (alternating 64mers and 112mers) –which represents a minimal HOR (Figure 2). Its ChIP/Input ratio (~13 with antibody against CENP-A) is elevated compared to the major satellite control (Figure 5), and compared to the 112mer, but much lower than that of the more common 120mer (~114). This elevated-but-low ChIP/Input ratio indicates that the 64mer is plausibly in the region flanking the centric core of one or more of the the repeat arrays that nucleate the kinetochores of the X and autosomes –but closer than the non-dimeric form of the 112mers. The lower copy number and lower diversity of the 64mer compared to both the 112mer and 120mer, is consistent with the hypothesis that this monomer –and the 176 bp dimer it forms with a 112mer– is younger. Because of the dimer’s longer length compared to the predominant 120mer, it may have the potential to eventually displace the 120mer –but only if it is repositioned (e.g., via ectopic gene conversion or mmBIR with template switching) to a region deep within the centric core of at least one chromosome (for the rationale of this hypothesis, see Rice 2019B).

The 120mer is by far the predominant monomer type (~89% of all monomers). Its much higher ChIP/Input ratio (~114; Figure 5), and the fact that FISH studies report this sequence to underly all active kinetochores on the X and autosomes (Wong and Rattner 1988), indicates that it is the principle repeat unit of *M. m. domesticus* centromeres. Analysis of contiguous monomers on Sanger tracers indicates that they are found predominantly within monotypic tandem arrays (Figure 2). Based on Illumina reads with low sequencing errors, the average divergence between randomly drawn 120mers was 5.9% (Figure 4). Using a sample of monomers from Sanger traces that are not side-by-side (on different traces), I get a similar value (5.75%; Supplemental Figure S7). The cluster analysis (neighbor-joining tree) of 120mer sequence variants based on sequence similarity (Figure 6) found two major sub-clusters: one with non-functional b-boxes and the other with predominantly functional b-boxes. The two subgroups consistently differ at four nucleotide positions: i) three of which are within the b-box sequence, ii) a fourth that is immediately adjacent to this b-box, and iii) two of which are predicted to completely suppress the binding of CENP-B. This high concentration of divergence within the b-box region that forms the major separation of 120mer clusters suggests that there may be functional significance to at least some of the polymorphism found among 120mers. The observation that more than half of the monomers carry mutated b-boxes that are predicted to fail to bind CENP-B (Supplemental Figure S4) suggests that non-binding monomers may have functional significance.

Sequence divergence among mouse 120mers is commonly described as low (e.g., Kipling et al. 1994; Kalitsis et al. 2006), and the estimates from this study are similar to previous estimates: 3.7% average sequence divergence from the 120mer consensus sequence (Figure S5) and an average of 5.9% divergence between random pairs of 120mers (Figure 4 and supplemental Supplemental Figure S7). These values might seem intuitively small, but comparisons to humans suggest they are not. In humans, average divergence of HOR units from the the consensus was reported to be only 1-2% on the × chromosome (Schueler et al. 2001) and divergence between pairs of the same monomer type within higher order repeats (located on chromosomes 8, 17 or X) was reported to be 0-3% (Schueler and Sullivan 2006). So sequence variation between mouse monomers is more than double that seen in humans when the same type of monomers within HORs are compared. It should be pointed out, however, that unlike the Western European house mouse that has only a single monomer type within its active centromeres on the X and autosomes, non-homologous humans chromosomes usually have different HORs and the monomers from non-homologous chromosomes can differ in sequence by as much as 35% (see Supplementary Table 1 in Rice 2019A for a genome-wide comparison of the consensus sequences of all human monomers at the active centromeres of a single genome).

An alternative way to quantify the sequence diversity at mouse centromeric monomers is to estimate the frequency spectrum of different monomer sequences, i.e., the plot of relative frequency vs. rank of abundance (Figure 7). This frequency spectrum demonstrates the extensive sequence diversity of these monomers: 1,923 unique sequences in a sample of 7,218 monomers (with low sequencing error rates) or an average of about about one new sequence for every fourth new monomer sampled. The frequency spectrum is characterized by an exceptionally long tail in which there were 827 variants with a frequency > 1 (Figure 7; note that low levels of sequencing error will generate almost exclusively novel sequence which will not contribute to this number). Even the most common sequence variant (which was the consensus sequence) was rare (a frequency of only 5.7%) and 80% of the monomer sequences had a frequency of less than 1%. So despite the moderate sequence diversity as measured by the average deviation of monomers from their consensus (3.7%) or average divergence between random pairs of monomers (5.9%), the frequency spectrum of monomer sequences demonstrates very high numerical diversity.

If monomer diversity is so exceptionally high, then why has it been overlooked in past studies? I think that the reason is that diversity can be defined to have three components and past studies have focused on only one of these. The first diversity component is the ‘richness’ component (or numerical diversity), which is the number of different sequences - the greater the richness the higher the diversity. The second diversity component is ‘evenness’, which is maximized when the distribution of variant frequencies is uniform and minimized when one or a few variants strongly predominate – the greater the evenness the higher the diversity. The third diversity component is sequence divergence among variants –the greater the average divergence the higher the diversity. Past studies have focused on only sequence divergence, but this component is not where most of the diversity of the Western European house mouse resides: it resides primarily in richness (numerical diversity) and evenness.

One might argue that the observed, unexpected pattern of exceptionally high monomer diversity might be some sort of an artifact of the the sliding window search algorithm used to find monomers. As a check on this possibility I used BLASTN on the Illumina data set to find monomers based on the the 120mer, 112mer, and 64mer consensus sequences. The high diversity of 120mers (high richness and evenness) were recapitulated in this alternative analysis, as were the relative frequencies of monomer size classes (Supplemental Figure S12).

The high level of sequence diversity found in the C57BL/6N inbred line might be due to the fact that: i) the same monomer is used at the centromeric repeat arrays of 20 different homozygous (inbred) chromosomes across the X and autosomes, and ii) different chromosome repeat arrays might drift to similar but non-identical sequences. For this reason, high global diversity (across all chromosomes) might occur despite low local diversity (within a single chromosomal repeat array or within subregions within these arrays). I was able to explore this possibility by examining monomers from long (containing ≥ 2 contiguous monomers), low-error-rate Sanger traces to compare sequence divergence at global and local scales. The first indication of high local diversity was the rare occurrence of side-by-side monomers with identical sequences –only 0.9% –and this value is similar to the global value (Supplemental Figure S6).

The generally accepted model for repeat array uniformity is the unequal crossing over model of Smith (1976). This model predicts a high level of sequence similarity at small spatial scales within a repeat array –yet side-by-side monomers rarely had the same sequence. A second way to compare local and global diversity is to compare sequence divergence between random monomers from the entire genome vs. side-by-side monomers (Supplemental Figure S7). This comparison indicated that nearly all global diversity is seen at the smallest possible spatial scale: side-by-side monomers (ave divergence = 5.32%) had 93% as much sequence divergence as random pairs of monomers (ave divergence = 5.75%).

To further explore global vs. local monomer sequence diversity, I made cluster diagrams from a sample of 25 5-monomer-long Sanger traces (Figure 6 and Supplemental Figure S9). The consistent pattern was for each 5mer to encompass most of the genome-wide, global sequence diversity –another indication of very high local diversity. These observations indicate that regional sequence divergence is not responsible for the high genome-wide diversity of monomers observed in the Western European house mouse.

The house mouse and humans have converged at their centromeres by independently evolving 17 bp b-boxes within their centromeres that bind CENP-B –a feature that increases connections between chromosomes and mitotic and meiotic spindles (Fachinetti et al. 2015). Yet despite this convergence, their centromeric repeat sequences are highly dissimilar: humans have complex multi-monomer HORs that differ among chromosomes while mice have a single active monomer that is shared by all the autosomes and X. Another distinguishing feature of mouse centromeres is extreme numerical diversity among the single monomer type found at the active centromeres: no sequence is common, tandem repeats of identical sequences are rare, and local diversity is nearly as high as global diversity. One process that might favor the high local sequence diversity of monomers observed in the Western European house mouse is repair of DSBs by the SSA repair pathway. This process deletes repeated sequences (usually one repeat unit per DSB) when they are located close to one another (Ozenberger and Roeder 1991). Another process is BIR repair of collapsed replication forks, which expands tandem rDNA repeat arrays (Kobayashi 2014). In the companion paper (Rice 2020), I explore the hypothesis that low sequence identity among nearby monomers is favored by molecular drive (Dover 1082) because it reduces the rate of deletions during repair of double strand breaks via SSA repair pathway –a repair mechanism that is expected to occur at elevated rates within the centromeres of telocentric chromosomes (Muraki and Murnane 2017).

Lastly, the strong level of sequence heterogeneity of monomers at a local level contrasted markedly with the high level of homogeneity of the of density monomers with and without functional b-boxes (those predicted to bind CENP-B and not bind this protein, respectively; Supplemental Figures S10 and S11). The pattern is similar to that found among the monomers at human centromeric HORs in which the primary sequence configuration is a dimer of different monomer types: one with and one without a functional b-box. This pattern in both humans and mice suggests that while binding of CENP-B strengthens kinetochores (Fachinetti et al. 2015), binding at too high a density may be suboptimal.

## Conclusions

The main findings from this quantification of centromeric monomer diversity within the genome of the Western European house mouse concern monomer length and sequence diversity. With respect to monomer length: i) there is substantial polymorphism, but only the 120mer predominates within the kinetochore-recruiting centric cores of the centromeric repeat arrays, ii) higher sequence diversity of the 112mer indicates that that it is the oldest monomer type and was plausibly replaced by the 120mer, iii) low sequence diversity of the 64mer indicates that it is the youngest monomer type and that this newer length variant (that is found as a 176mer dimer with 112mers) may have the potential to become the predominant repeat unit in the future. The major findings for the sequence diversity of the 120mers (that recruit the kinetochores of the mouse chromosomes) are: i) despite modest sequence divergence from the consensus sequence, the monomers have an exceptionally diverse frequency spectrum (high richness) with no common sequence variants and many hundreds of types of low-frequency variants (high evenness), ii) identical sequences are rare in side-by-side monomers, iii) sequence divergence between side-by-side monomers is nearly as great as divergence genome-wide, iv) sequence divergence within small ‘neighborhoods’ of five contiguous monomers typically spans nearly all the diversity found genome wide, and v) the densities of monomers with and without functional b-boxes are far more uniform than predicted by chance. The companion paper explores a hypothesis to explain these patterns.

## Acknowledgments

I thank two undergraduates working in my laboratory (Roselyn Wu and San Ha Lee) for assistance during the exploratory stages of this research. I also thank Kathryn Schoenrock for copy-editing assistance.

## Supplemental Figures

**Supplemental Figure S1.**
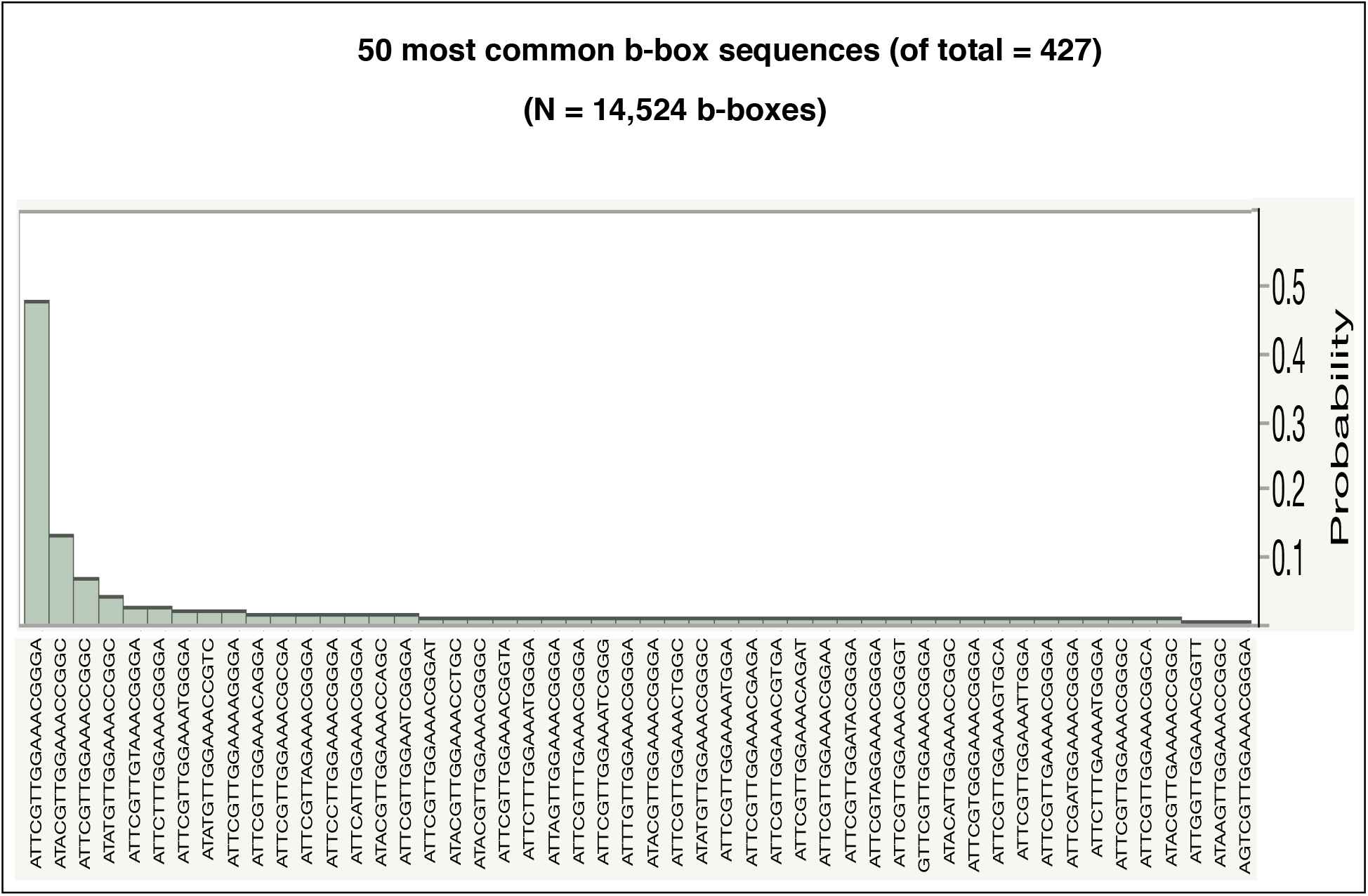
The 50 most common b-box sequences observed when the consensus sequences from the 64mer, 112mer, and 120mer monomers were used as query sequences with BLASTN using 150 bp Illumina reads (first 42,211,594 lines of SRA file *SRR6339177;* strain C57BL/6J; Wang et al. 2018). This generated 14,524 b-box sequences which were clustered into 427 different sequence categories.

**Supplemental Figure S2.**
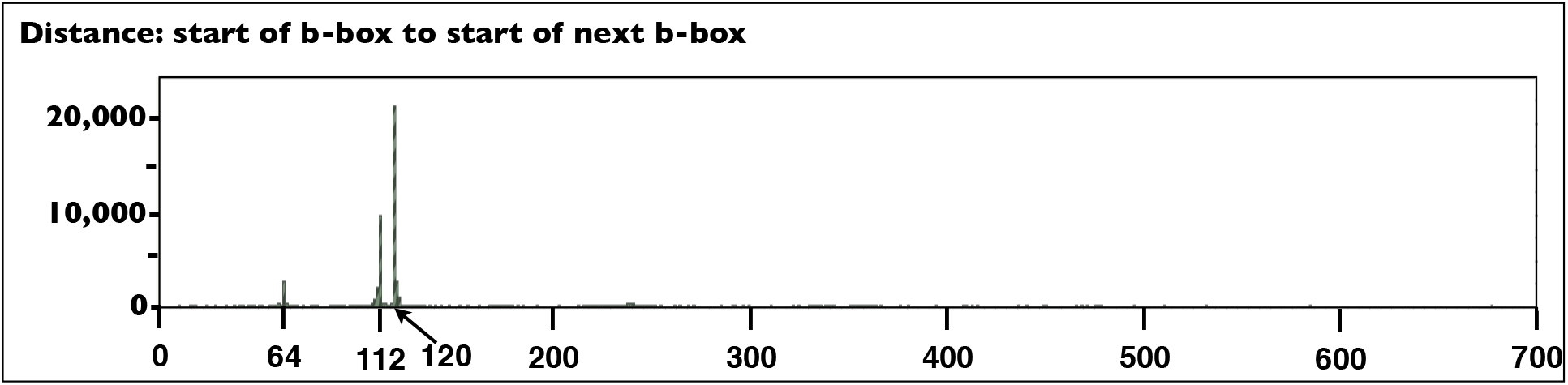
A plot of abundance vs. distance between the start of a b-box and the start of the next b-box along Sanger traces. The b-boxes were detected by a 17 bp sliding-window that was run along the length of each Sanger trace. A b-box was scored as present whenever the 17 bp window matched the 17 bp consensus b-box sequence with ≤ 4 bp mismatches (or matched one of seven 5 bp deviations that were found to be present at non-trivial frequency in b-boxes). There were three major peaks at 64, 112, and 120 bp.

**Supplemental Figure S3.**
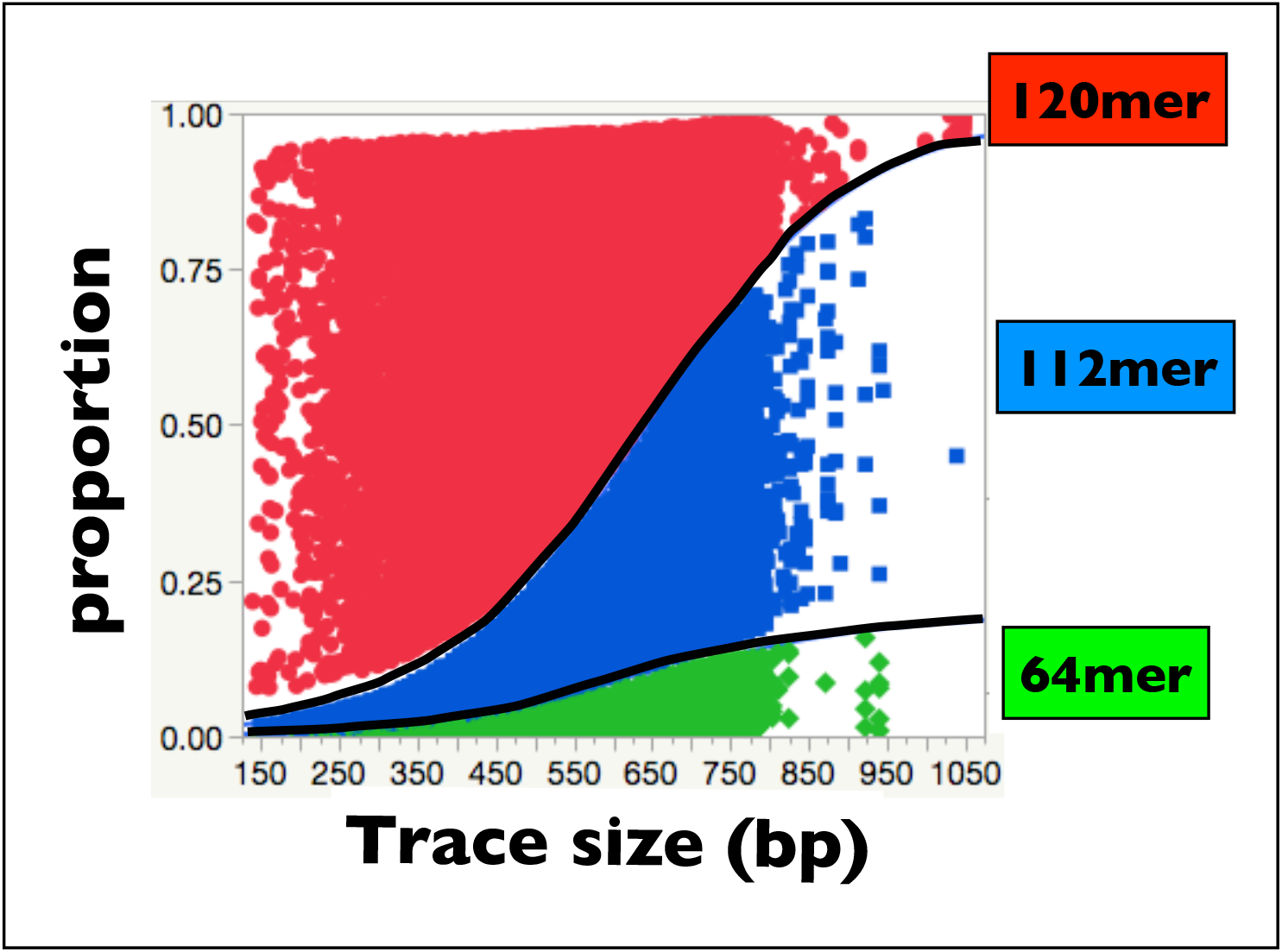
Logistic regression of the categorical variable ‘monomer type’ (64mer, 112mer, or 120mer) on Sanger trace length. There is a strong influence of trace length on the relative frequency of the three monomer types –with 120mers frequency declining with increasing trace length. N = 41,539 monomers from traces containing at least one monomer from the Mouse Genome Project trace files 1-49 (ftp://ftp.ncbi.nih.gov/pub/TraceDB/mus_musculus/).

**Supplemental Figure S4.**
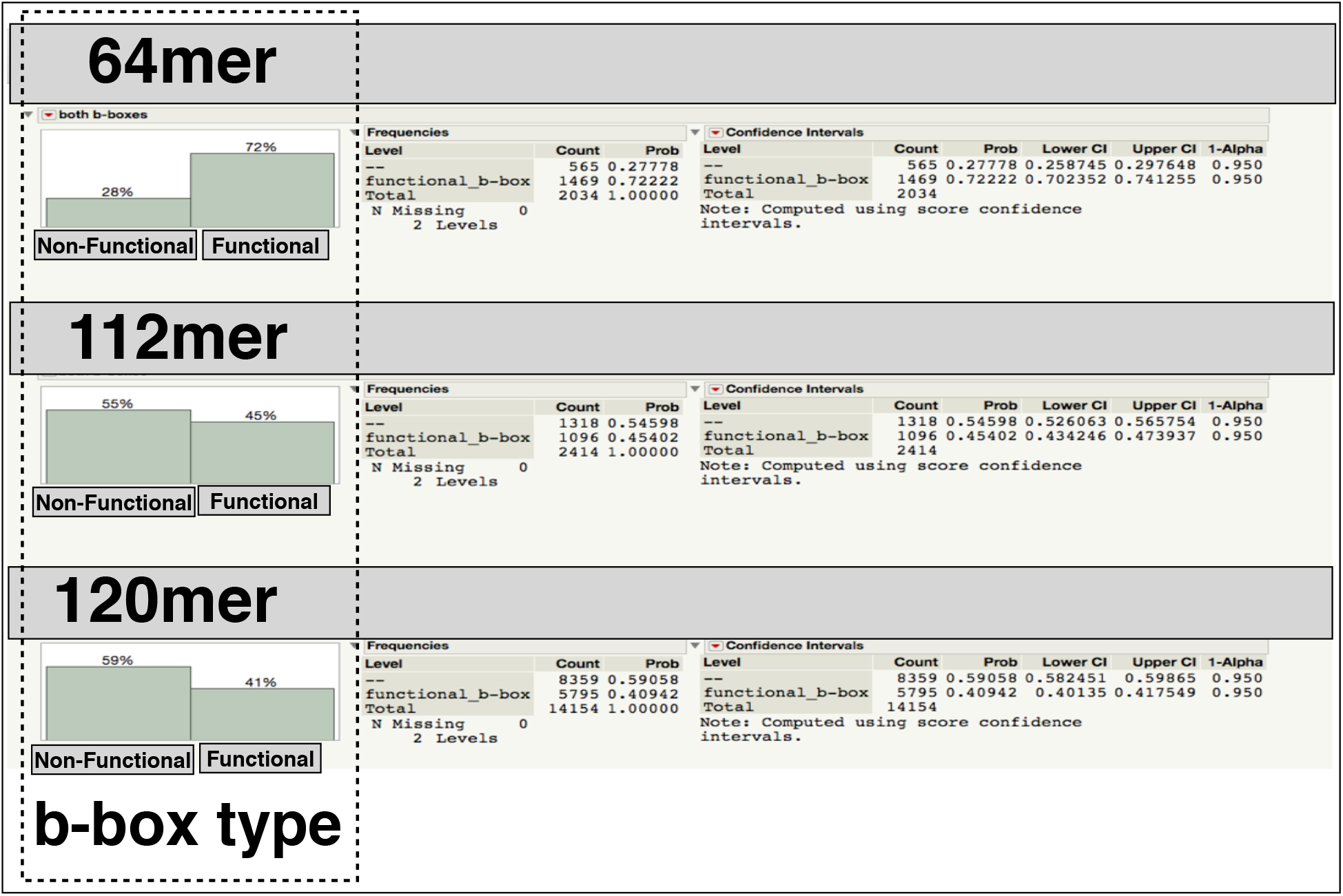
Comparing 64mer, 112mer and 120mers monomers for the proportions with and without function b-boxes, i.e., those predicted to bind –and not to bind– CENP-B. All monomers used in the comparisons have an average accuracy per bp of 99.9% or higher.

**Supplemental Figure S5.**
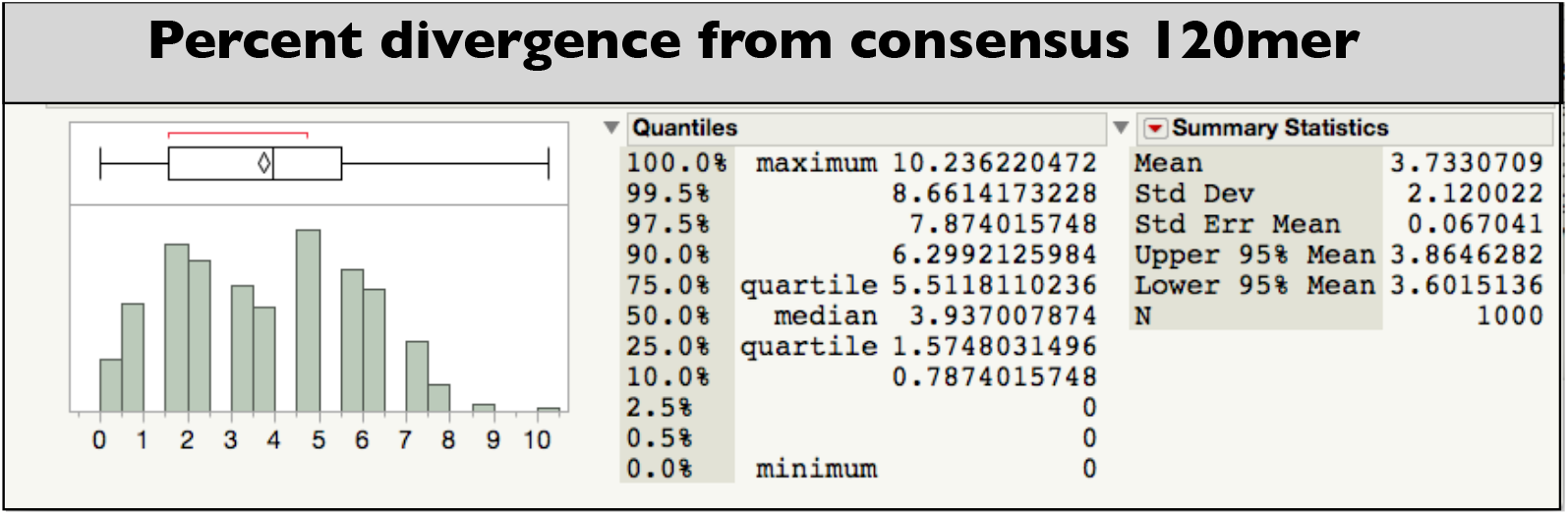
The percent divergence from the 120mer consensus sequence of a sample of monomers with sequence accuracy per bp of 99.9% or higher.

**Supplemental Figure S6.**
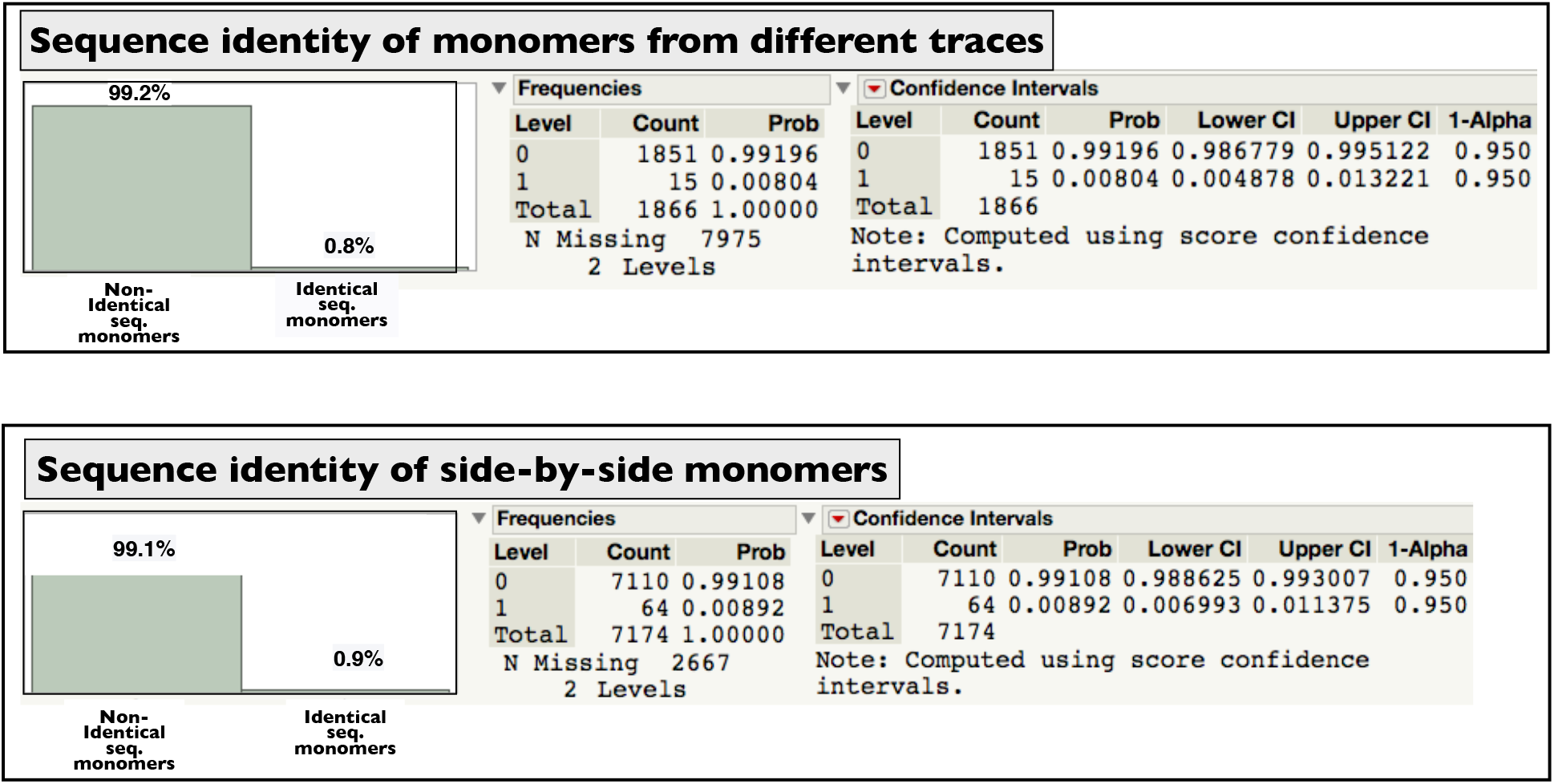
The proportion of pairs of monomers with identical sequence (sequence identity) for random pairs of monomers (taken from different Sanger traces) or side-by-side monomers on the same Sanger trace. All monomers in the sample have an average sequence accuracy per bp of 99.9% or higher.

**Supplemental Figure S7.**
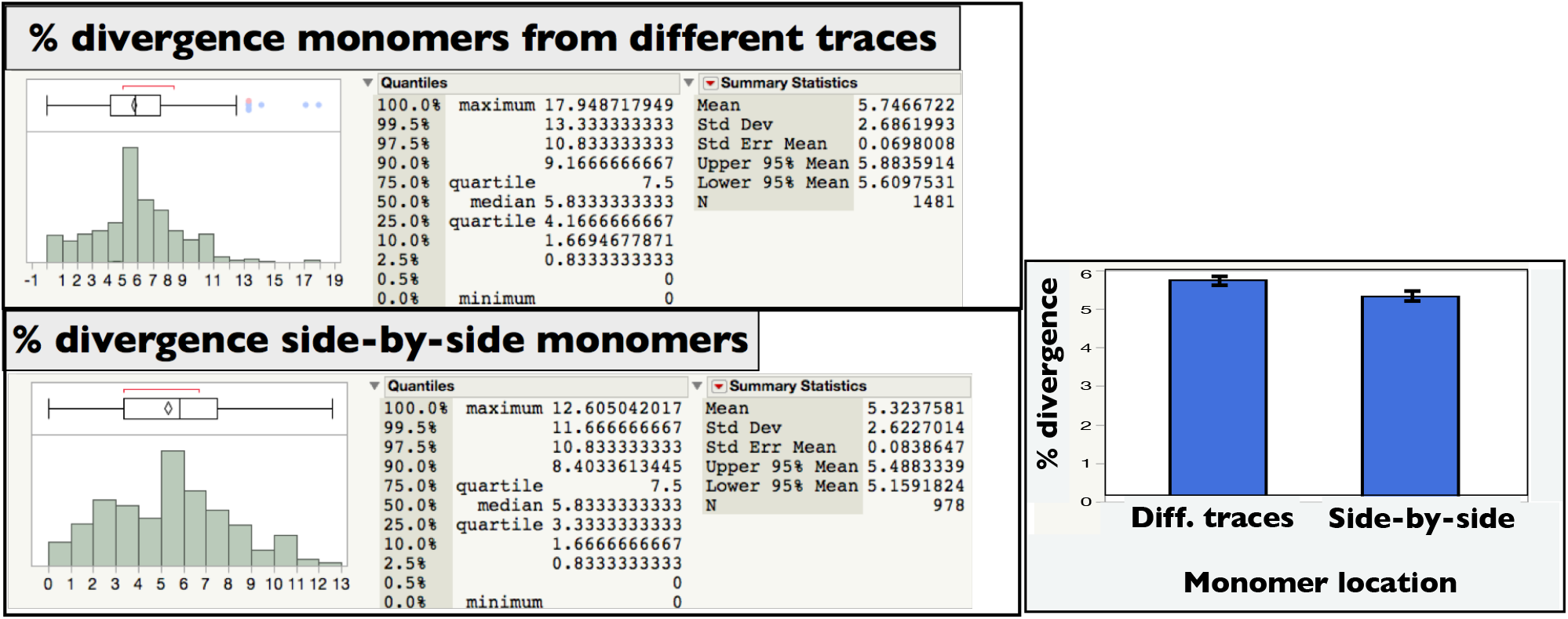
The percent divergence (percent miss-matching base pairs across the length of a pair of monomers) for random pairs of monomers (taken from different Sanger traces) or side-by-side monomers on the same Sanger trace. All monomers in the sample have an average sequence accuracy per bp of 99.9% or higher. Note that although the % divergence estimates are similar, the 95% CIs for the mean do not overlap.

**Supplemental Figure S8.**
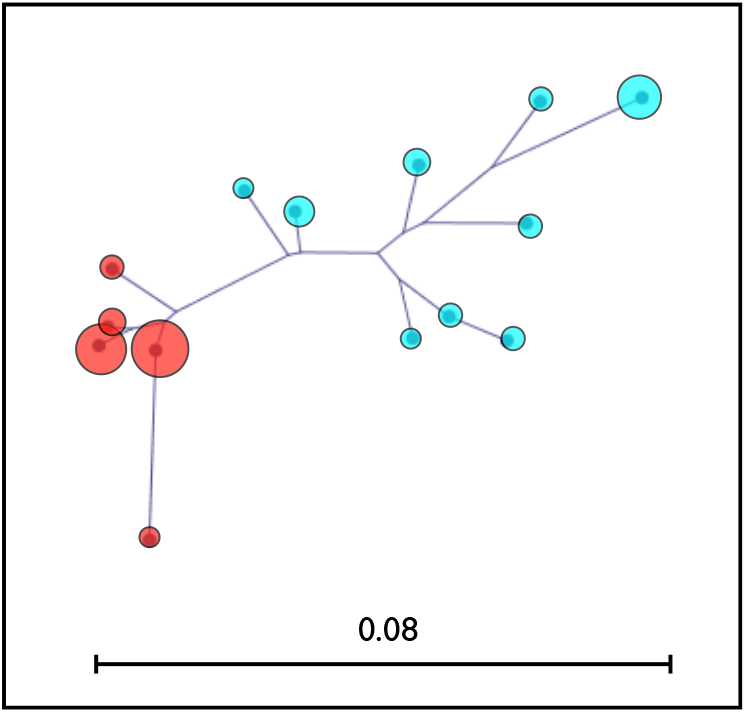
A cluster diagram for the 13 most common 120mer monomer sequences. The sequence for the 18th most common was also included to have the clustering cover all the major groups shown in Figure 6. See Figure 6 for fuller description of the nodes.

**Supplemental Figure S9.**
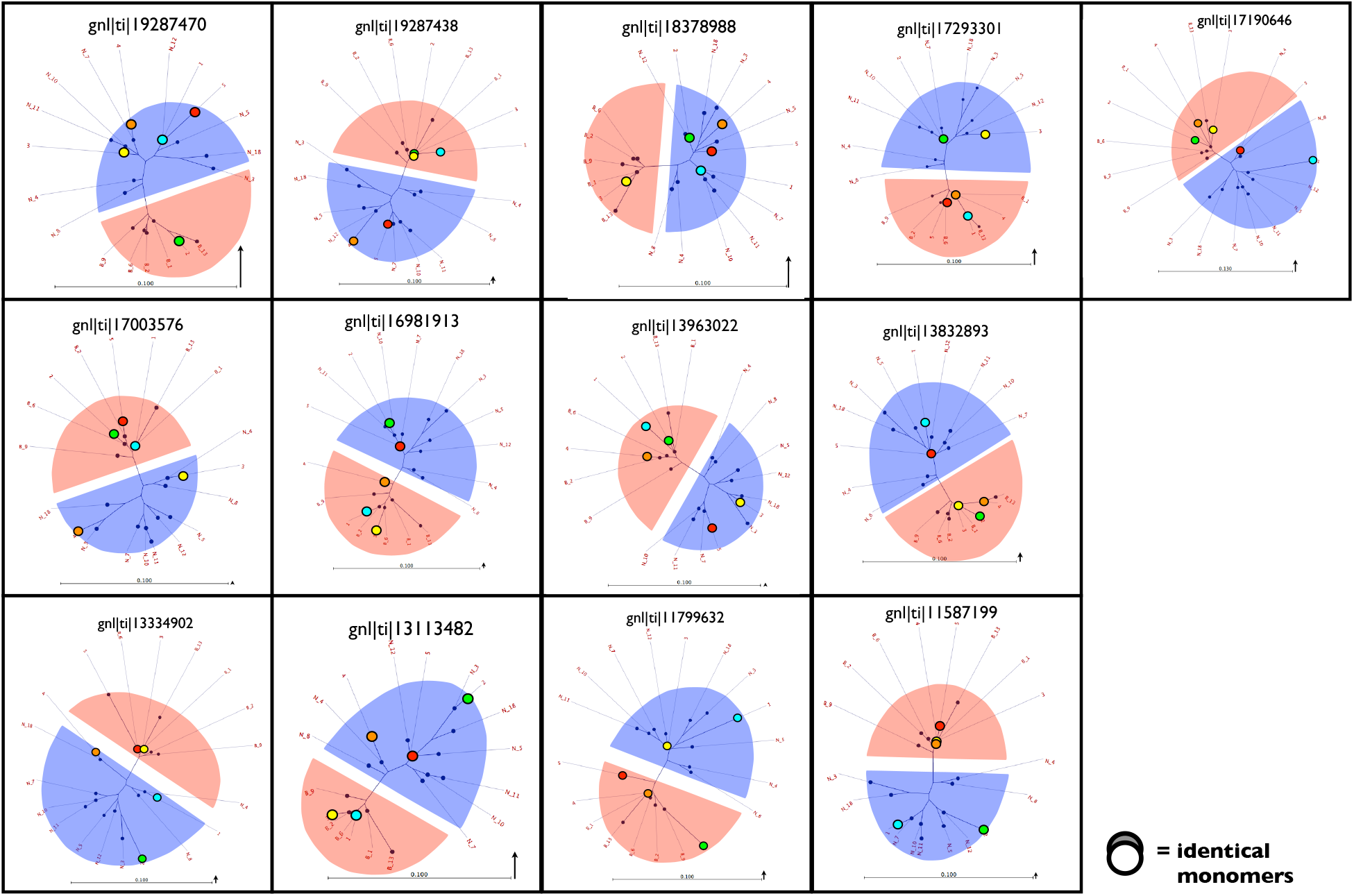
An extension of the cluster diagrams shown in Figure 8 that includes the remaining 13 (out of a total of 25) Sanger traces with higher sequencing error rates. See Figure 8 for fuller description.

**Supplemental Figure S10.**
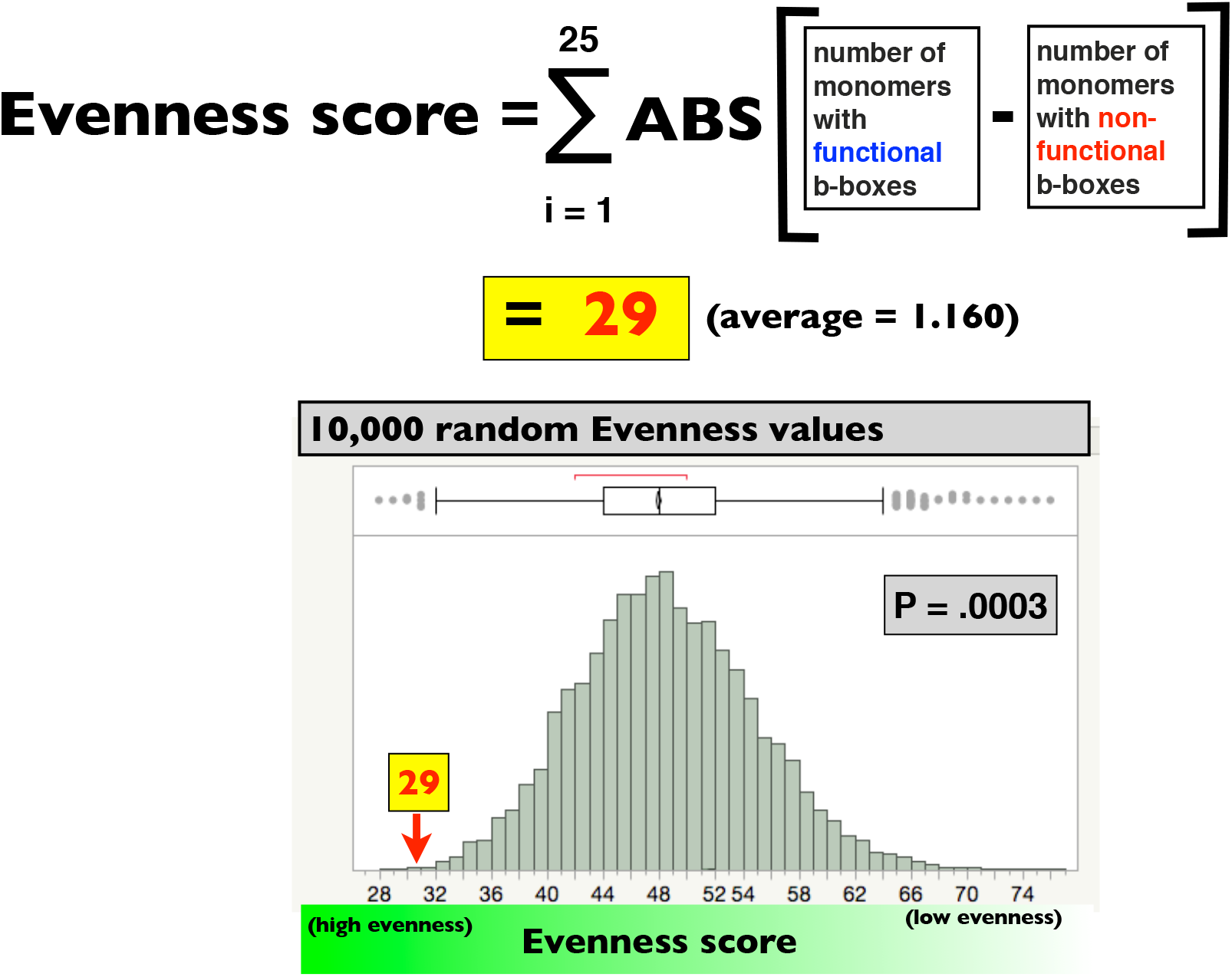
A randomization test to examine the statistical significance of the observed high level of evenness in the frequency of monomers with and without functional b-boxes (17 bp sequence binding CENP-B). In the sample of 25 long Sanger traces (containing five contiguous 120mers) the evenness score was 25 (top of figure) and 10,000 randomly generated evenness cores were rarely this small (P = 0.0003).

**Supplemental Figure S11.**
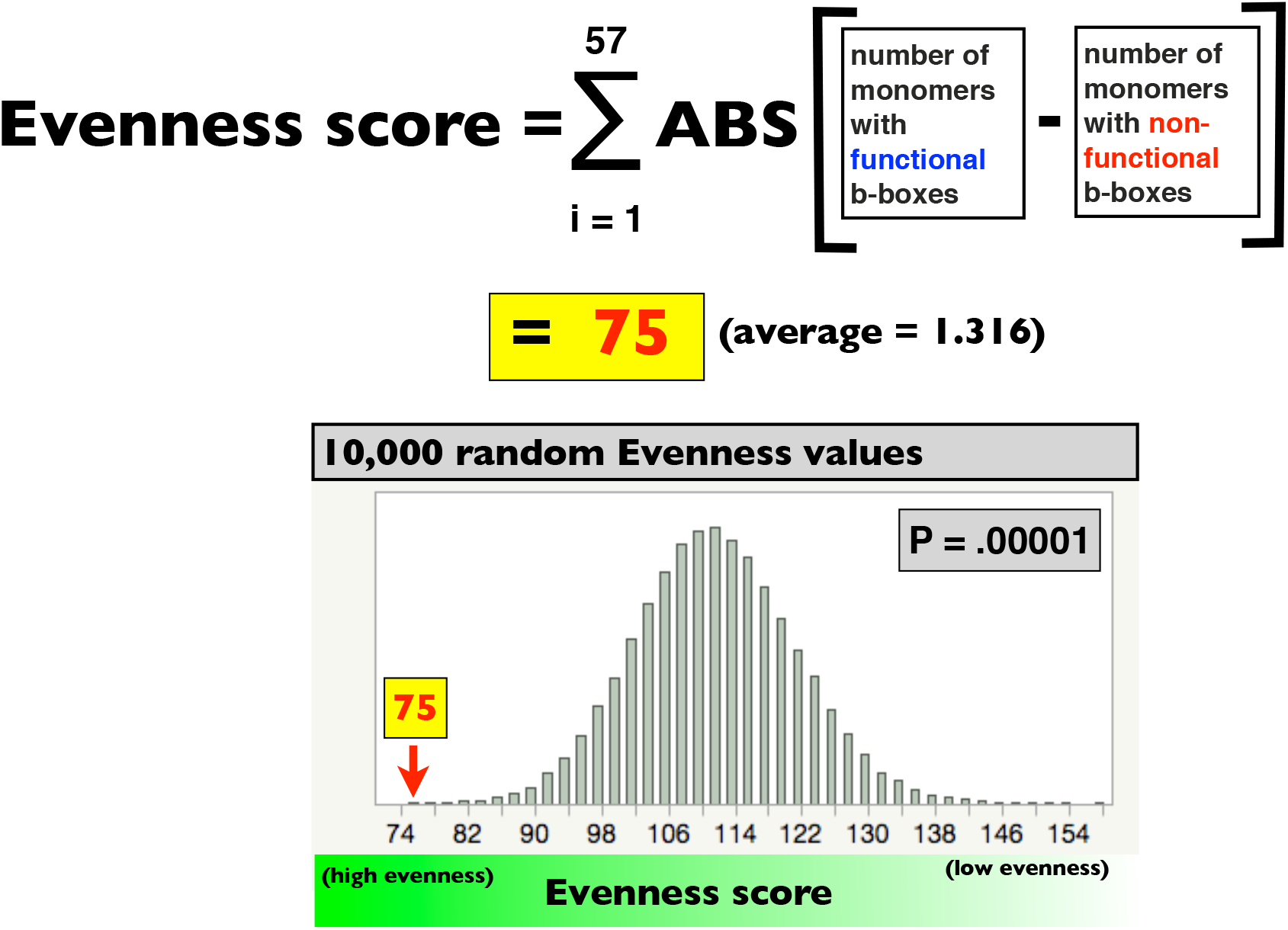
Extension of Supplemental Figure S10 to a larger sample of Sanger traces with 5 contiguous 120mer monomers. In the larger sample of 57 long Sanger traces (containing five contiguous 120mers) the evenness score was 75 (top of figure) and 100,000 randomly generated evenness scores were rarely this small (P = 0.00001).

**Supplemental Figure S12.**
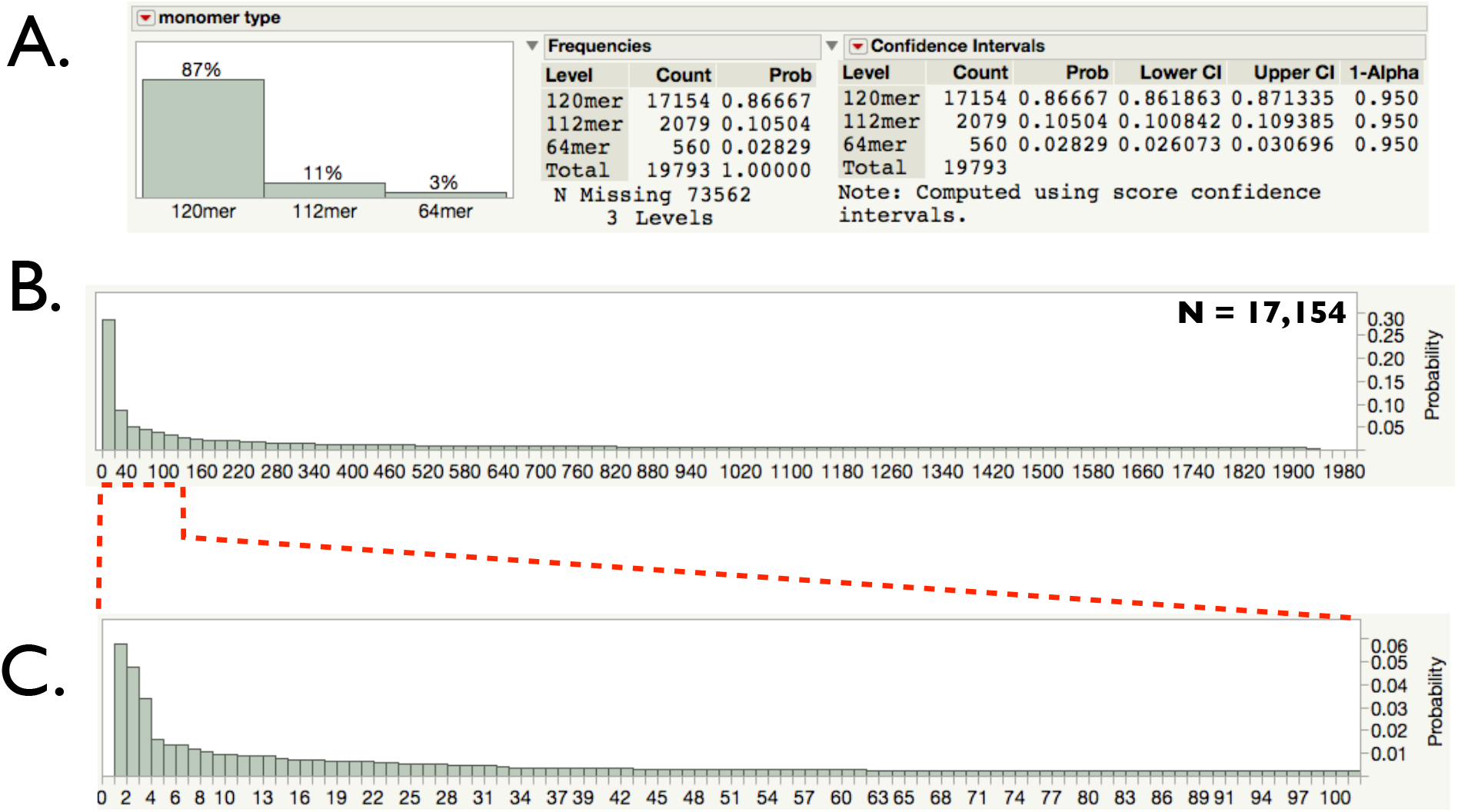
Mouse centromeric monomers from BLASTN. The consensus 120mer, 12mer and 64mer (Figure 1) were used to BLAST the 150 bp Illumina reads of Wang et al. (2018) [SRA file SRR6339177] and extract monomer sequences. To reduce the influence of sequencing error, only reads having all bp calls with ≥ 99.9% accuracy were used. **A.** The distribution of monomer size class variants (from a sample of 19,793 low-sequencing error monomer sequences) closely matched that of the sliding window analysis (Figure 2). **B-C.** The frequency spectrum of 17,154 120mers (plot of the rank of each monomer sequence’s frequency vs. its rank in frequency) closely matched that from the sliding window analysis (Figure 7) – confirming high richness diversity and high evenness diversity among 120mers.

